# Translation efficiency is maintained at elevated temperature in *E. coli*

**DOI:** 10.1101/109264

**Authors:** Gareth J Morgan, David H Burkhardt, Jeffery W Kelly, Evan T Powers

## Abstract

Cellular protein levels are dictated by the balance between gene transcription, mRNA translation and protein degradation, among other factors. Cells must manage their proteomes during stress; one way in which they may do so, in principle, is by differential translation. We used ribosome profiling to directly monitor translation in *E. coli* at 30 °C and investigate how this changes after 10-20 minutes of heat shock at 42 °C. Translation is controlled by the interplay of several RNA hybridization processes, which are expected to be temperature sensitive. However, translation efficiencies are robustly maintained after thermal heat shock and after mimicking the heat shock response transcriptional program at 30 °C. Several gene-specific parameters correlated with translation efficiency, including predicted mRNA structure and whether a gene is cotranslationally translocated into the inner membrane. Genome-wide predictions of the temperature dependence of mRNA structure suggest that relatively few genes show a melting transition between 30 °C and 42 °C, consistent with our observations. A linear model with five parameters can predict 33% of the variation in translation efficiency between genes, which may be useful in interpreting transcriptome data.

## INTRODUCTION

The regulation of the rate of protein synthesis is not completely understood. Cells continuously alter protein levels and stoichiometry in order to maintain a correctly-balanced proteome. Chaperones, proteases and specialized folding factors ensure that proteins can reach their native state without becoming trapped in misfolded or aggregated states (1, 2). The rate of synthesis of proteins needs to be tuned so that the capacity of this “protein homeostasis network” is not exceeded (3). In this work, we aimed to determine how *E. coli* alter their translation in response to heat shock by using ribosome profiling (4, 5) to measure genome-wide translation rates. These data represent an important step towards understanding how bacteria strike a balance between synthesis, folding and degradation of proteins, and will facilitate the development of computational models of the *E. coli* protein homeostasis network (3).

An archetypal stress on proteomes is heat shock, abruptly raising cells’ temperature so that proteins unfold and aggregate. Rapid heating in bacteria triggers the heat shock response, a transcriptional program characterized by increased expression of chaperones and proteases to enhance protein folding and suppress aggregation. The canonical heat shock genes are transcribed using the alternate RNA polymerase sigma factor, σ^H^, encoded by the *rpoH* gene (6, 7). σ^H^ activity is tightly regulated at several levels, including enhanced translation at elevated temperatures, and binding by its primary downstream targets (8–10). Production of new, misfolding-prone proteins is a major source of protein aggregation and toxicity during heat shock, so we reasoned that differential translational control may be important in reducing the concentrations of heat-labile proteins.

Control of translation plays an important role in the expression of many genes (5, 11–15), reviewed in (16). Over a population of cells, mRNA levels generally correlate with protein levels, but there are large differences between individual proteins (12, 14, 17). A clear example in prokaryotes is the differential translation of individual genes from polycistronic transcripts (12, 15). Control of translation has been intensely studied, but there is little consensus on the relative roles of different factors, and no way to predict how well-translated a particular sequence will be (16). Translation rate is determined by a combination of initiation and elongation rates. Translational initiation is slower than elongation, so initiation is rate-limiting for the translation of most genes (18).

Ribosome profiling by deep sequencing measures the distribution of translating ribosomes within a cell’s transcriptome, and hence determines how often a particular mRNA is translated (4, 5, 12). A key parameter is translational efficiency (TE), the rate of protein production per mRNA, which is equivalent to the number of ribosomes that translate a particular mRNA molecule. This is a more direct measurement of translation than is possible by measuring protein levels, and in combination with total mRNA measurements can assess genome-wide TE. Ribosome profiling can report on protein synthesis rates, which in turn correlate strongly with protein abundances in rapidly-dividing cells such as *E. coli* (12). Surprisingly, differences in TE measured by ribosome profiling correlate weakly, if at all, with sequence-specific factors that affect translation of individual genes, and the determinants of differential TE across transcriptomes remain unknown (12, 16, 19). Recent work suggests that ORF-wide mRNA structure is the primary determinant of TE differences in the *E. coli* genome (20).

Here, we use ribosome profiling to directly quantify the relationship between mRNA abundance and ribosome occupancy at 30 °C and under heat shock conditions (42 °C) in the well-studied model organism, *E. coli* K12 MG1655. We find that TE and patterns of ribosome footprints for all measured genes are very similar between 30 °C and after 10 or 20 minutes of heat shock at 42 °C despite widespread changes in transcription and translation levels. mRNA structure seems to play a significant role in determining TE. RNA stability predictions suggest that few mRNAs undergo structural transitions in the temperature range studied. Unrelated to our original hypothesis, we did observe one striking and unexpected correlation: A distinctly lower TE for inner membrane proteins under both normal and heat shock conditions, which we hypothesize is linked to cotranslational export from the cytosol.

## RESULTS

*Translation efficiency varies across the E. coli genome*—We sequenced total RNA and ribosome footprints from replicate *E. coli* K12 MG1655 cultures growing exponentially in rich defined media (Table 1). We tested the hypothesis that heat shock would alter translation by sequencing libraries from bacteria growing at 30 °C and after 10 and 20 minutes of growth at 42 °C. Any changes in translation that we observed could be directly caused by temperature-dependent RNA hybridization, or by downstream effects of genes expressed at high temperature. To differentiate between these possibilities, we also compared bacteria expressing either wild-type or I54N σ^H^ protein from a pBAD plasmid, which mimics the heat shock transcriptional program, to bacteria containing an empty pBAD vector at 30 °C (21). σ^H^, encoded by the *rpoH* gene, is the RNA polymerase sigma factor responsible for the transcription of the canonical heat shock proteins such as the chaperones DnaK and GroEL. The activity of σ^H^ protein is repressed by factors including DnaK, but this repression is alleviated by the I54N mutation (10, 22).

**Table 1:**
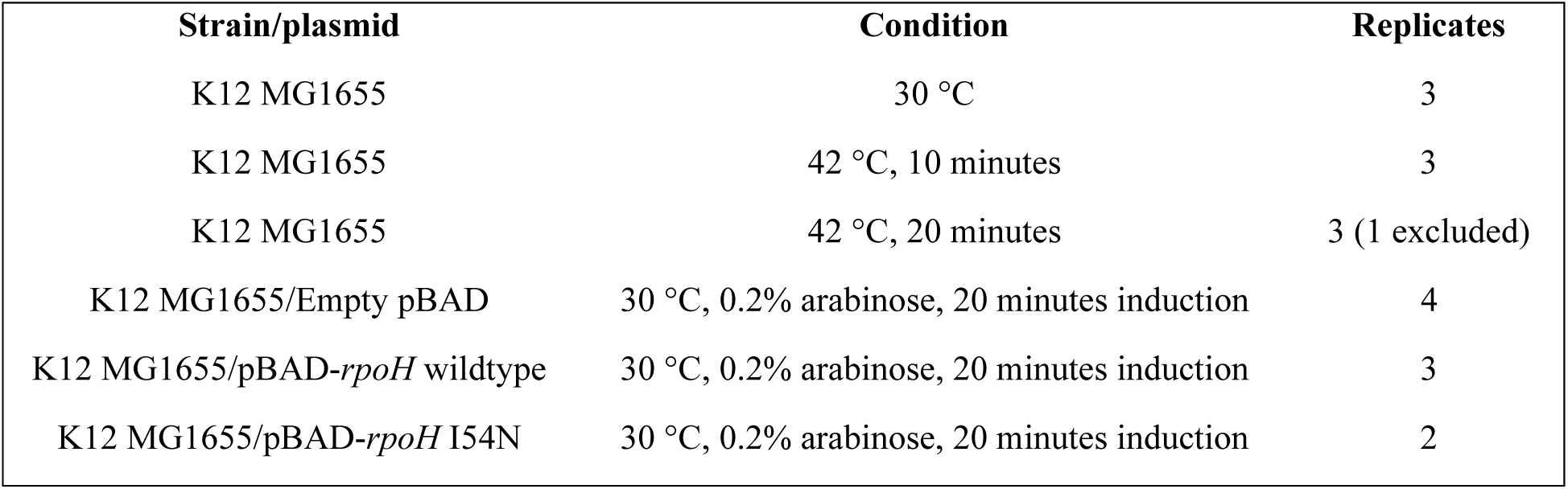
Ribosome profiling experiments. See also Table S1.

| Strain/plasmid | Condition | Replicates |
| --- | --- | --- |
| K12 MG1655 | 30 °C | 3 |
| K12 MG1655 | 42 °C, 10 minutes | 3 |
| K12 MG1655 | 42 °C, 20 minutes | 3 (1 excluded) |
| K12 MG1655/Empty pBAD | 30 °C, 0.2% arabinose, 20 minutes induction | 4 |
| K12 MG1655/pBAD-*rpoH* wildtype | 30 °C, 0.2% arabinose, 20 minutes induction | 3 |
| K12 MG1655/pBAD-*rpoH* I54N | 30 °C, 0.2% arabinose, 20 minutes induction | 2 |

For each sample, mRNA and ribosome footprint cDNA libraries were prepared and sequenced. The number of reads per gene was calculated and normalized with EdgeR (23) to give mRNA and footprint counts per kilobase million (CPKM) for each gene. Footprint reads were adjusted to remove the influence of the elevated ribosome density seen at the beginning of genes (12). We focus on protein coding genes without unusual translational events such as frameshifting. For each gene, and for each replicate, we calculated translation efficiency (TE), the ratio of footprint CPKM to mRNA CPKM. Homologous genes such as *tufA* and *tufB*, which encode elongation factor EF-Tu, were considered as a single gene and their counts summed. We included data from *E. coli* grown at 37 °C in similar rich defined media obtained by Li and co-workers (12) in our analysis for comparison. Normalized gene read counts for each library are shown in Table S1. The footprint reads from one replicate of the 20 minute heat shock condition differ substantially from those of the other replicates (Table S1), suggesting a problem with library construction, so those data were excluded from our analysis.

Under all conditions, global transcription and translation patterns are similar, with CPKM values varying over 1000-fold between genes (Fig. 1A and 1B). Footprint levels correlate well with mRNA levels (Fig. 1, R^2^ = 0.80 for 30 °C log-transformed data), indicating that transcript level is a primary determinant of translation rate. However, translation efficiency varies by more than 100-fold across the genome (Fig. 1C and 1D), even within individual operons. Data for an extreme example, the *yobF*-*cspC* operon, is highlighted in Fig. 1A. Translation levels for measurable proteins correlate with those observed previously (5, 12), and with proteomic abundance measurements (Fig. 1E and 1F), (24, 25).

**Fig. 1:**
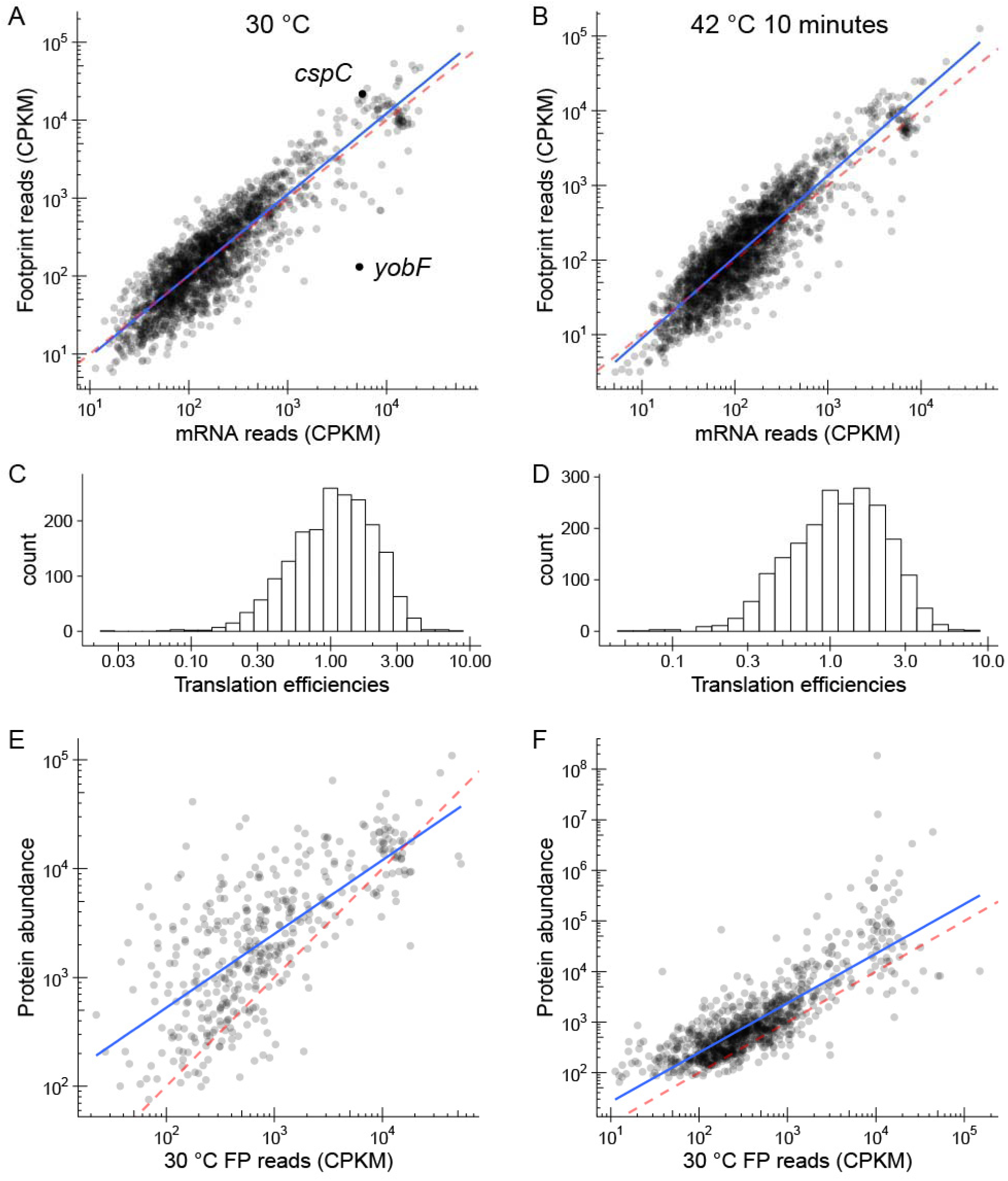
Translation varies across the genome, and within operons. Ribosome footprint levels are strongly correlated with mRNA levels at both 30 °C (A) and after 10 minutes heat shock at 42 °C (B). Each point represents a gene. The blue lines show linear regression fits to the log-transformed data, while the red lines are 1:1 ratios. Although mRNA level is the primary determinant of translation level, the ratio of footprint to mRNA reads varies by over 100-fold across the genome, as illustrated by the *yobF-cspC* operon, highlighted in A. Histograms of translation efficiencies at 30 °C (C) and after 10 minutes heat shock at 42 °C (D) show similar distributions of translation. (E) Ribosome footprints at 30°C correlate with absolute protein abundance measurements by mass spectrometry (25) (F) Ribosome footprints at 30 °C correlate with protein abundance measurements by mass spectrometry (24). CPKM = counts per kilobase million.

*Translation efficiency is maintained at elevated temperature—*Because translation initiation is controlled by the interplay between different RNA hybridization events, we reasoned that temperature should differentially affect the TE of different genes. However, the measured TEs of genes do not significantly change between 30 °C and 42 °C (Fig. 2), despite changes in absolute translation levels between conditions (Fig. 3). We looked for altered ratios of footprint and mRNA counts using EdgeR (23) and anota (26); neither method identified genes with significantly altered TE (not shown).

**Fig. 2:**
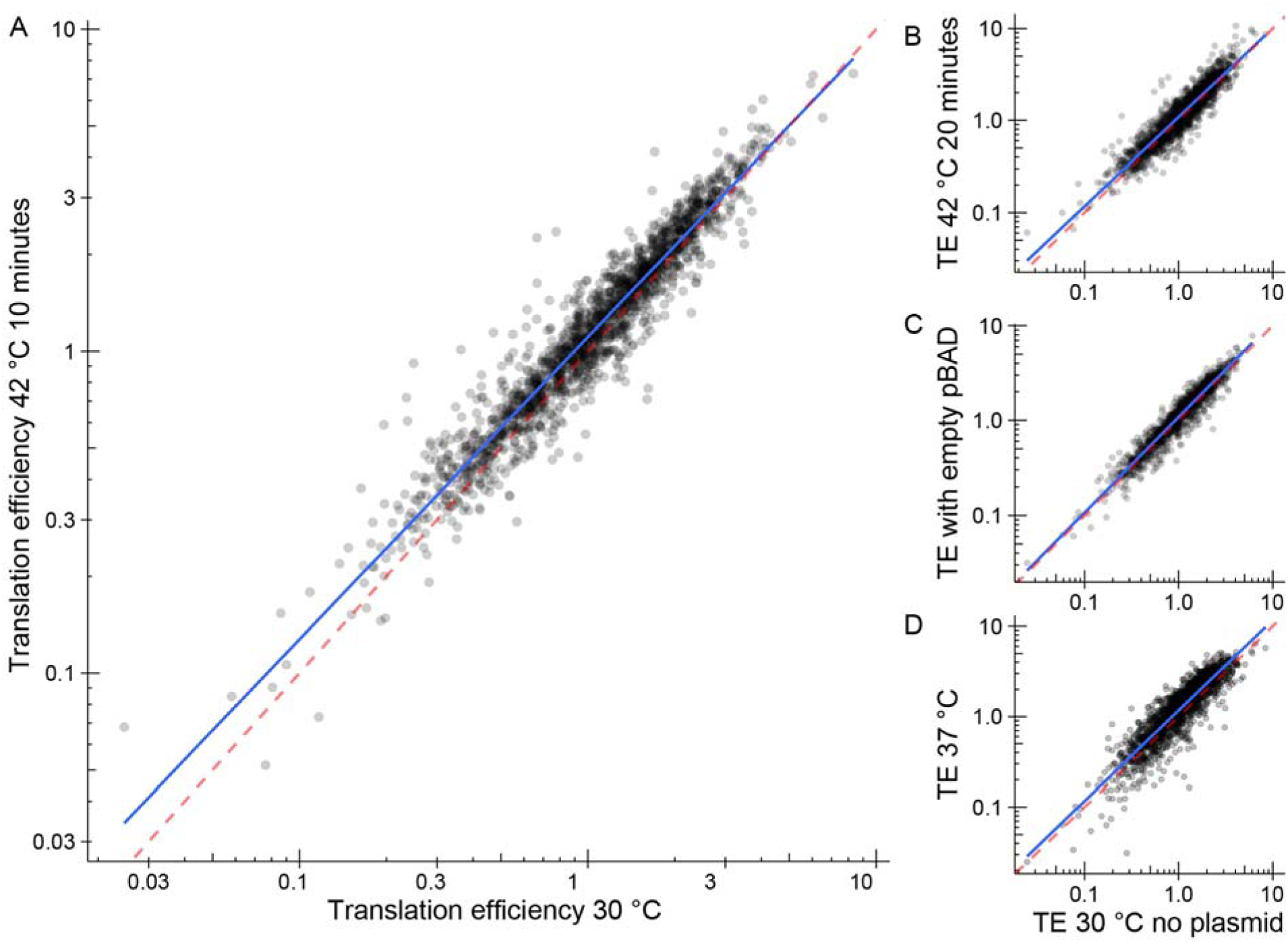
Translational efficiency is maintained during heat shock. Plots of translational efficiency per gene at 30 °C versus (A) after 10 minutes at 42 °C; (B) after 20 minutes at 42 °C; (C) with an empty pBAD plasmid; and (D) compared to data at 37 °C from Li et al. (12).

**Fig. 3:**
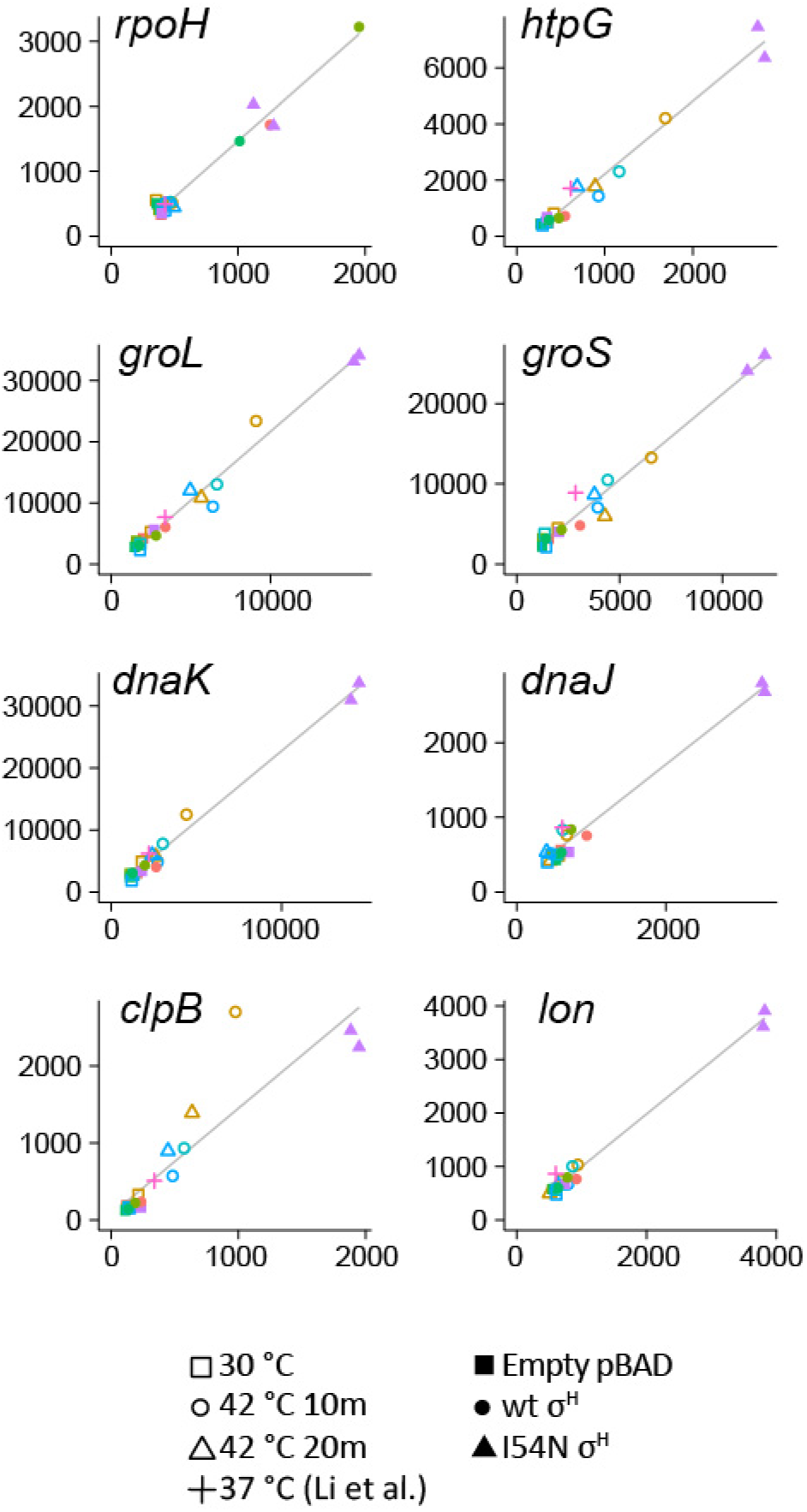
Translational efficiency of genes is independent of heat shock and expression level. Footprint versus mRNA levels are shown for several well-characterized genes whose expression changes upon heat shock, under different conditions. Shapes represent different conditions, and colors represent biological replicates grown on separate days. Linear fits are shown in gray. The ratio of footprint to mRNA reads is similar under all conditions tested, independent of the expression level of the gene. Data from Li et al. (12) at 37 °C is included for comparison.

Fig. 3 shows plots of footprint versus mRNA counts for selected canonical heat shock genes under different conditions. The strong induction of genes such as the chaperonin subunit *groL* indicates that the cells are responding to heat stress by transcribing genes from the σ^H^ regulon. Since the data in each case have a strong tendency to fall on a line, it is clear that TE (the slopes of the lines) is maintained at different temperatures and widely varying expression levels. Individual genes within polycistronic operons often have differing ribosome densities, indicative of differential translation (5, 12, 15). The patterns of translation within individual operons, which here are very similar to those described previously (12), are also maintained at different temperatures, as exemplified by the *rpsM* and *rpsP* operons (Fig. 4).

**Fig. 4:**
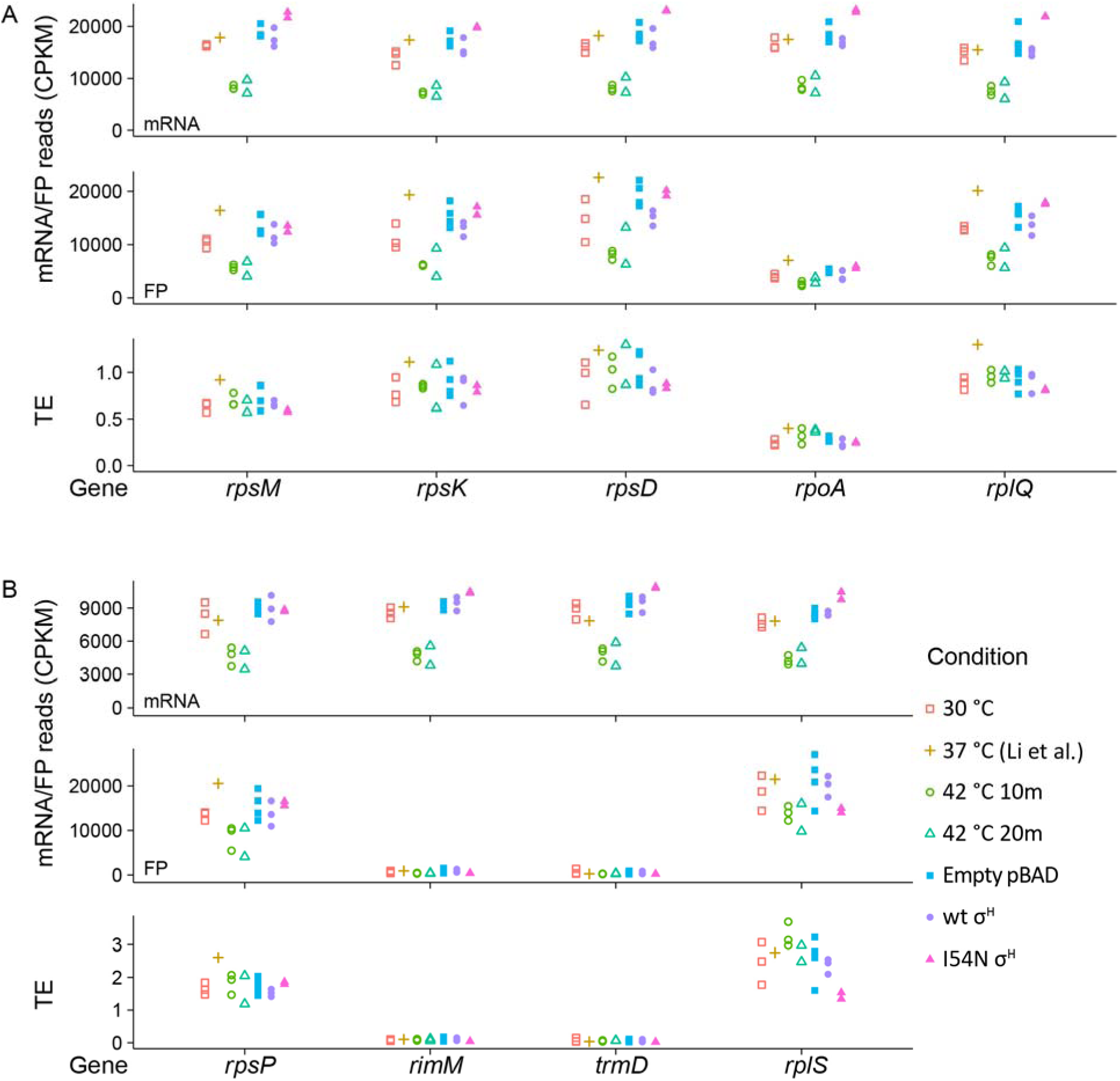
Differential translation within the ribosomal operons *rpsM* (A) and *rpsP* (B) is maintained at different temperatures and mRNA levels. The non-ribosomal genes *rpoA*, *rimM* and *trmD* have been shown to be differentially translated from the ribosomal proteins on these polycistronic operons. Data from Li et al. (12) at 37 °C is included for comparison.

*Ribosome footprint patterns are maintained at elevated temperature*—Ribosomes are distributed unevenly across mRNA molecules, showing distinct patterns in different genes. Translation slows or pauses when ribosomes interact with sequences that resemble Shine-Dalgarno motifs (27). Since this interaction is driven by RNA hybridization, we looked for changes in the patterns of ribosome footprints across the genome at different temperatures. Fig. 5A shows the ribosome profiles of the *dnaJ* gene, whose expression increases upon heat shock. The patterns of ribosomal density are very similar between conditions and replicates, indicating that ribosome pausing is not affected by heat shock. The similarity between profiles can be simply measured by correlation. The pairwise correlation coefficients for replicates of the same condition, and different conditions within the same replicate for *dnaJ* are shown in Fig. 5B. There is a stronger correlation between data measured at different temperatures than between replicates of the same temperature. This observation is likely due to the replicates having been grown on different days, whereas the six samples for each replicate (mRNA and footprints for 30 °C and for 10 and 20 minutes at 42 °C) were processed and sequenced together. This pattern is repeated across the genome for genes that are well expressed in all samples (Fig. 5C).

**Fig. 5:**
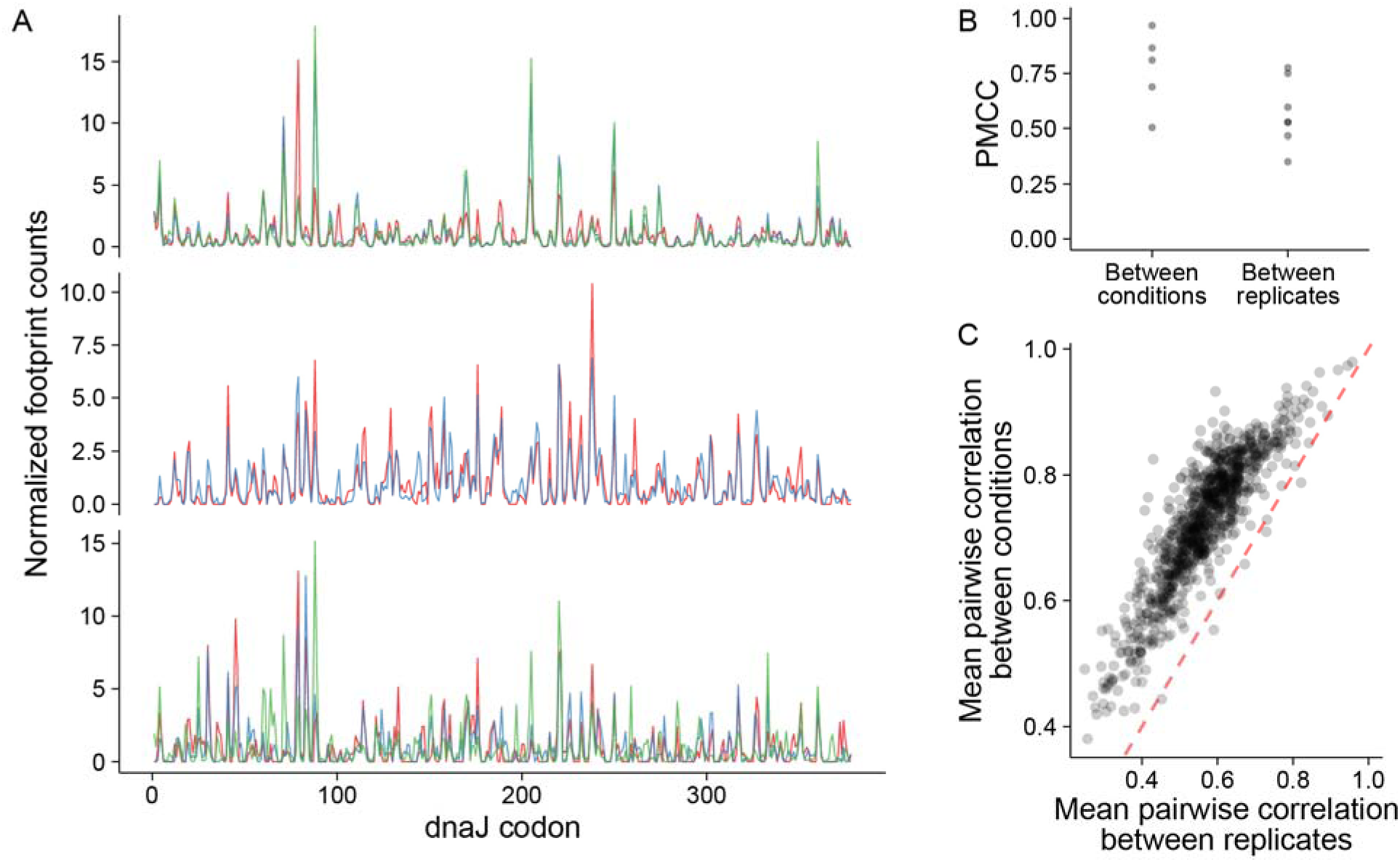
Patterns of footprints do not change between conditions. (A) Distribution of normalized footprint reads across the *dnaJ* gene for the three biological replicates. Colors represent different conditions. (B) Pairwise correlations between conditions for each replicate, and between replicates within each condition of footprint distributions for the *dnaJ* gene. (C) Mean pairwise correlations as for panel B for 939 genes with at least 64 footprint reads in each dataset. The red line shows a 1:1 correspondence.

*Activation of σ*^*H*^ *is controlled post-translationally*—The heat shock sigma factor, σ^H^, is regulated at several levels: translation of its mRNA is limited by secondary structure, and the protein is rapidly bound by DnaK and the signal recognition particle (SRP) and delivered to the membrane to be degraded by the FtsH protease (8, 10, 28, 29). As the downstream targets of σ^H^ are produced, they compete with RNA polymerase apoenzyme for binding to σ^H^, which turns off the transcriptional response. We use transcription of σ^H^ regulon genes as a measure of σ^H^ activity, and thereby distinguish between translational and post-translational regulation. Fig. 6 shows σ^H^ activity — measured by transcription of the *groL* and *dnaK* genes, which encode two major heat shock chaperones — as a function of its translation. At 30 °C, the activity of wild-type σ^H^ is only weakly dependent on its translation level, either from the chromosome or a plasmid. Translation of *rpoH* mRNA is similar at 30 °C and after 10 minutes at 42 °C. However, transcription of both *groL* and *dnaK* increases (hollow circles in Fig. 6) to a much greater extent than is seen following the overexpression of wild-type σ^H^ at 30 °C (solid circles in Fig. 6). After 20 minutes at 42 °C, translation of *rpoH* remains at a similar level, but σ^H^ protein activity decreases (hollow triangles in Fig. 6). Finally, direct overexpression of I54N σ^H^ results in a large transcriptional activity (solid triangles in Fig. 6). This pattern is consistent with post-translational repression of σ^H^ by DnaK and SRP being the primary means of control at ambient or mild heat shock conditions.

**Fig. 6:**
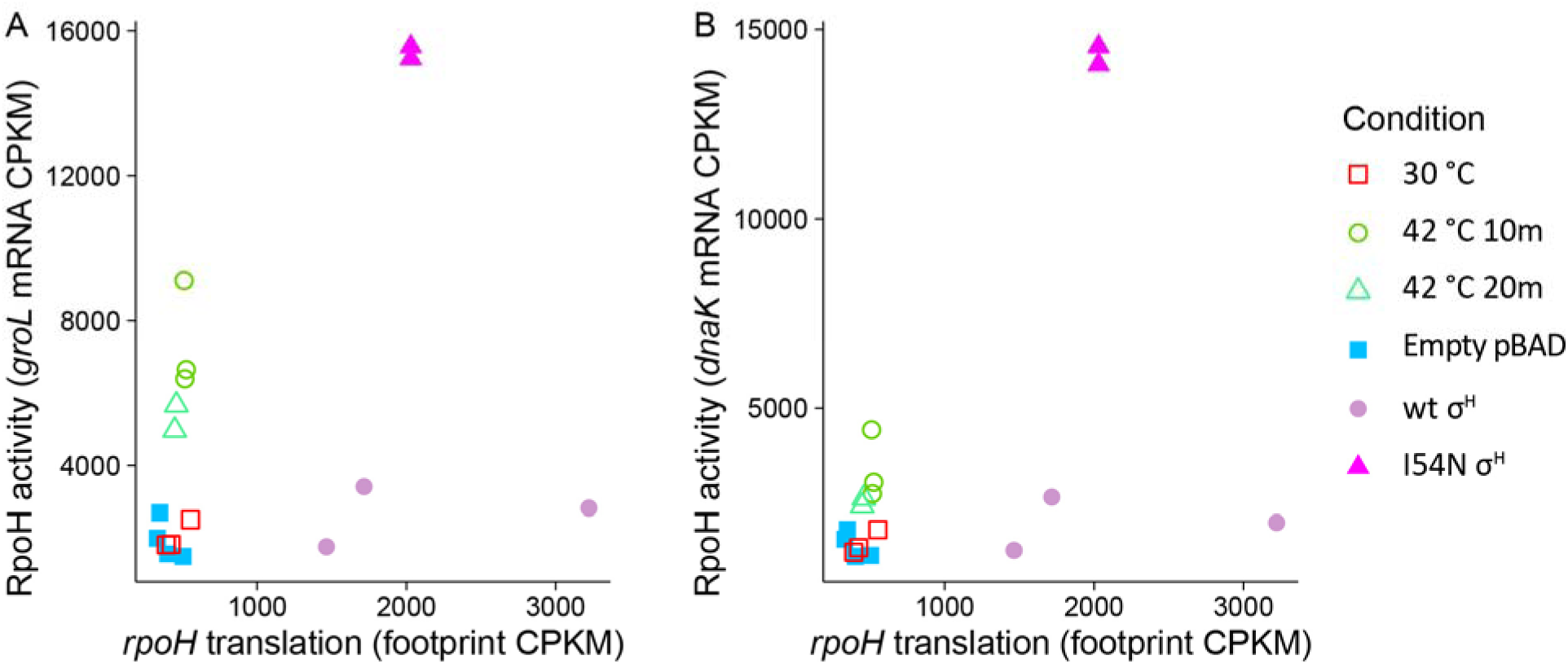
Activity of σ^H^ is primarily controlled post-translationally. The σ^H^ sigma factor, encoded by the rpoH gene, is not translationally upregulated by the heat shock treatment (footprint CPKMs of 456 ± 86 at 30 °C and 518 ± 7.4 at 42 °C after 10 minutes). However, its transcriptional targets, represented by the canonical heat shock proteins GroEL (A) and DnaK (B) are upregulated by heat. Overexpression of mutant, but not wild-type σH results in increased target transcription. These observations are consistent with the release of σ^H^ from post-translational control by DnaK/DnaJ and SRP upon heat shock.

*E. coli* has two other known RNA thermometers. Similarly to *rpoH*, the translation of the small heat shock protein IbpA is known to be controlled by RNA secondary structure that occludes its ribosome binding site at low temperature (30). We do not observe enough reads for *ibpA* to be able to reliably assess its translation, but its low expression in itself suggests that its translation is not activated. The cold shock protein CspA’s mRNA contains a motif that activates its translation at low temperature (31). However, *cspA* is strongly translated (TE of 4.6 ± 1.1 at 30 °C and 4.5 ± 0.77 after 10 minutes at 42 °C), suggesting that this mechanism does not prevent translation under these conditions.

*Translation from the open reading frame of ssrA increases during heat shock*—*E. coli* have several mechanisms to rescue ribosomes that have stalled on an mRNA molecule, the best-understood of which is the tmRNA/*ssrA* system (32, 33). The tmRNA molecule, encoded by the *ssrA* gene, binds to ribosomes with a stalled nascent peptide, which may be caused by an mRNA lacking a stop codon. The tmRNA molecule releases the ribosome from the mRNA and encodes for the translation of a short peptide tag which directs the resulting peptide for degradation by the ClpXP protease. FP counts from the ORF portion of the *ssrA* gene increase following heat shock (Fig. 7). This indicates an increase in ribosome stalling during heat shock, possibly caused by an increased frequency of mRNA fragmentation or translational frameshifting.

**Fig. 7:**
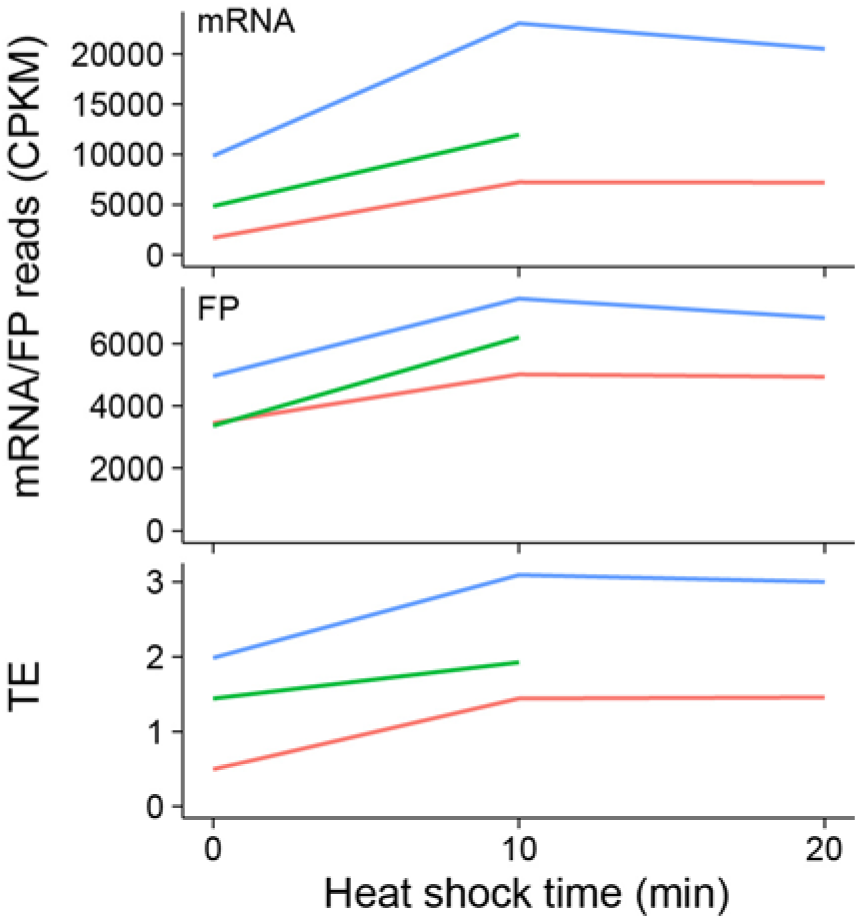
Translation from the coding region of the *ssrA* gene increases during heat shock. Both mRNA and footprint levels increase after 10 minutes of heat shock and remain elevated after 20 minutes of heat shock. Colored lines represent replicate cultures.

*Translation efficiency differences between genes are partly determined by ORF-wide mRNA structure*—Since TE does not change significantly between conditions, but varies widely between genes (Figs. 1 and 2), it must be regulated by the cell. There are several known gene-specific factors that can influence translation in *E. coli*. Translation initiation is thought to be rate limiting in most cases, and this is controlled by a combination of ribosome binding to mRNA, mRNA secondary structure and codon use. We therefore examined the relationship between these metrics and TE, expecting that factors known to influence translation rate would correlate with differences in TE. We describe the results for the 30 °C dataset, but other conditions have similar patterns.

Recent work suggests that *E. coli* mRNA molecules are organized into ORF-wide structures, and that the extent of these structures determines TE (20). In agreement with this, we found a negative correlation between TE and the predicted stability of an ORF’s mRNA sequence, corrected for gene length (Fig. 8A, R^2^ = 0.15). A weaker correlation exists between TE and the tRNA adaptation index (34), a measure of codon use (Fig. 8B, R^2^ = 0.14). However, factors that influence translation initiation are less well correlated with TE. Similarly to previous studies, we see no correlation between calculated ribosome binding site strength and TE. The effect of a gene’s start codon is smaller than we expected. Fig. 8C shows the distribution of translation efficiencies for genes as a function of their start codon. While genes with non-AUG start codons are, on average, less well-translated than those with AUG start codons, the effect on TE is small (mean TEs of 1.31 for AUG vs 0.988 for non-AUG, p < 10^-10^, Welch’s t-test) and there is no significant difference in TE between UUG and GUG codons.

**Fig. 8:**
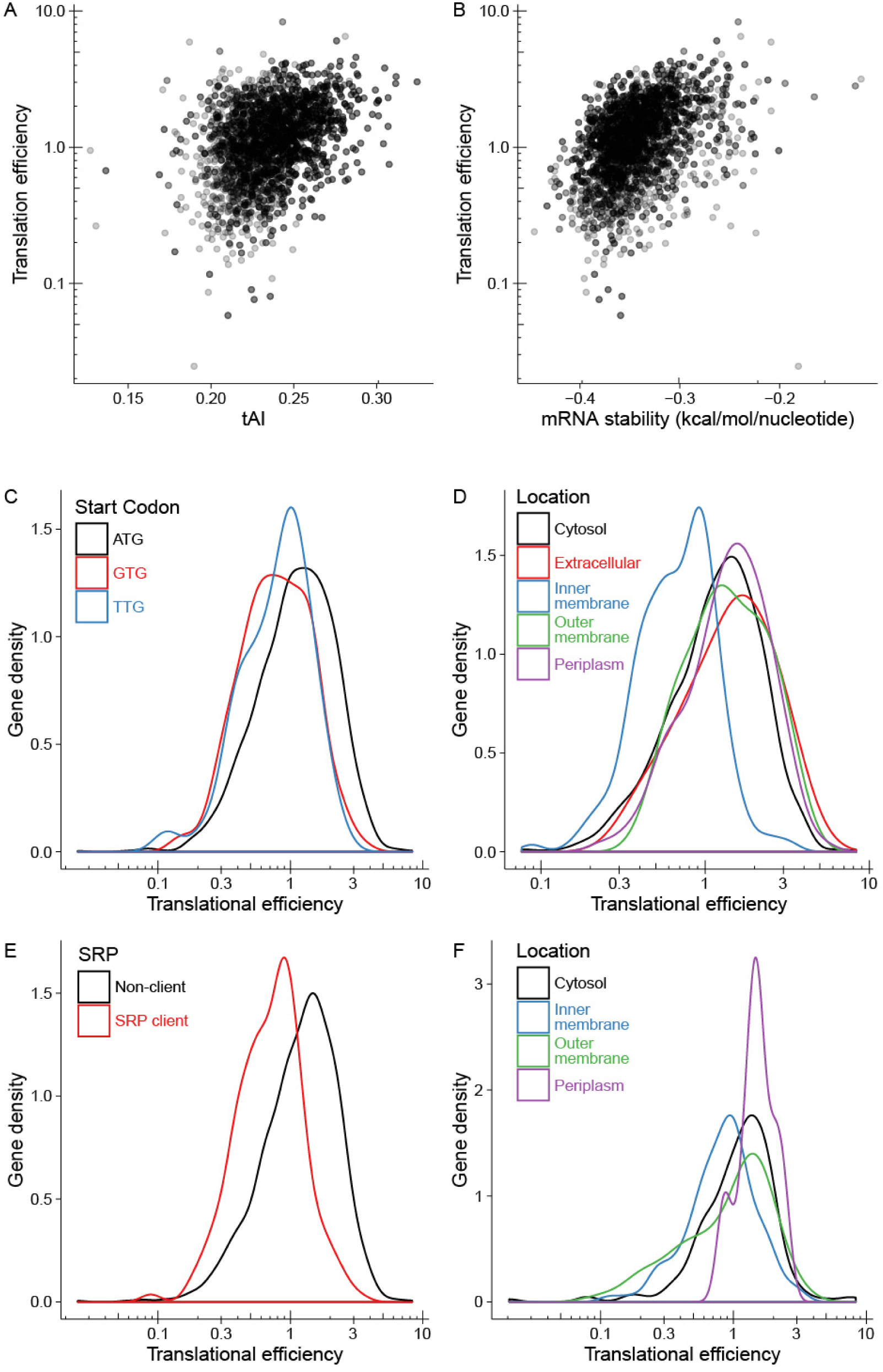
ORF structure, tRNA availability, start codon identity and cellular location affect translation efficiency. (A) Correlation between tRNA adaptation index and TE. (B) Correlation between ORF mRNA secondary structure stability (normalized for gene length) and TE. (C) Non-AUG start codons are disfavored in but not excluded from highly translated genes. Smoothed density histogram of translation efficiencies for the three most highly-represented start codons. Curves are scaled so that their integrals are equal. (D) Inner membrane proteins are less highly translated than other proteins. Density histogram of translation efficiency for genes identified as either residing in the cytosol, inner membrane, periplasm, outer membrane, or secreted from the cell. (E) Clients of the SRP pathway are less efficiently translated than other proteins. Density histogram of translation efficiency for genes identified as SRP clients by Schibich and coworkers (37). (F) *Caulobacter crescentus* inner membrane proteins are less translated than proteins in other subcellular locations. Ribosome profiling data on proteins with known localization in *C. crescentus* was taken from Schrader and coworkers (38). Gene translation efficiencies were calculated for *C. crescentus* growing in rich media.

*Inner membrane proteins are significantly less well translated than other classes of proteins*—All *E. coli* proteins are translated in the cytosol but many are co- or post-translationally exported to the periplasm, inner or outer membranes, or secreted from the cell (35). Distributions of translational efficiency as a function of protein location, taken from the consensus locations defined in Diaz-Mejia et al (36), are shown in Fig. 8D. Inner membrane proteins have significantly lower TE than proteins destined for any other cellular compartment (mean TEs of 0.801 for inner membrane proteins and 1.40 for others, p<10^-15^, Welch’s t-test). Notably, outer membrane proteins and periplasmic proteins, which are also exported from the cytosol, have very similar patterns of translational efficiency to cytosolic proteins. The TE of a protein could be influenced by its mode of translocation. Most inner membrane proteins are cotranslationally translocated into the membrane via the SecYEG or YidC translocons, in a process dependent on SRP (35, 37). SRP clients have been identified using selective ribosomal profiling and are predominantly inner membrane proteins (37). The TE of SRP clients is significantly lower than that of non-clients, supporting the hypothesis that some feature of SRP-dependent synthesis affects translational efficiency (Fig. 8E). We analyzed ribosomal profiling data on the Gram negative bacteria *Caulobacter crescentus* (38) and found a similar reduction in TE for inner membrane proteins to that seen in *E. coli* (Fig. 8F), suggesting that this feature is conserved at least in these closely-related bacteria.

*Temperature-dependent mRNA structural transitions are infrequent between 30 °C and 42 °C*—The correlation between ORF structure and TE raises the question of whether general thermal unfolding of mRNA structures might be expected to increase TE. Our data suggest that this is not a major effect at the temperatures studied here. To further investigate this, we calculated the temperature dependence of mRNA stability for all protein coding genes using the RNAheat program from the Vienna package (39). The resulting melting curves (also known as thermograms) show peaks at temperatures where unfolding events occur. Example melting curves are shown in Fig. 9A. The midpoint temperatures of these events (11,399 in all genes, 5,104 in genes with temperature-dependent TE data) are shown as a histogram in Fig 9B. Most genes have similar predicted melting curves, with a large transition around 90 °C. Only a minority of transitions occur between 30 °C and 42 °C, and many of these transitions have small enthalpies compared to the total folding free energy of the gene’s mRNA. The 241 genes with transitions having excess heat capacities estimated to be > 0.01 kcal/mol/nucleotide in the 30 °C to 42 °C range are highlighted in Fig. 9C. Only 95 of 1813 genes with measured TEs at 30 °C and 42 °C are both weakly translated at 30 °C (TE < 1) and are predicted to have a structural transition between 30 °C and 42 °C. This is in broad agreement with the maintenance of TE for the majority of genes at these temperatures. This relative lack of thermal transitions suggests that mRNA structure is generally selected to be insensitive to temperature changes in this range, and that TE is therefore buffered against temperature change by mRNA structure. However, the RNA structure calculations are not well constrained for such long molecules, and these results may not be representative of the mRNA structural ensembles in living cells.

**Fig. 9:**
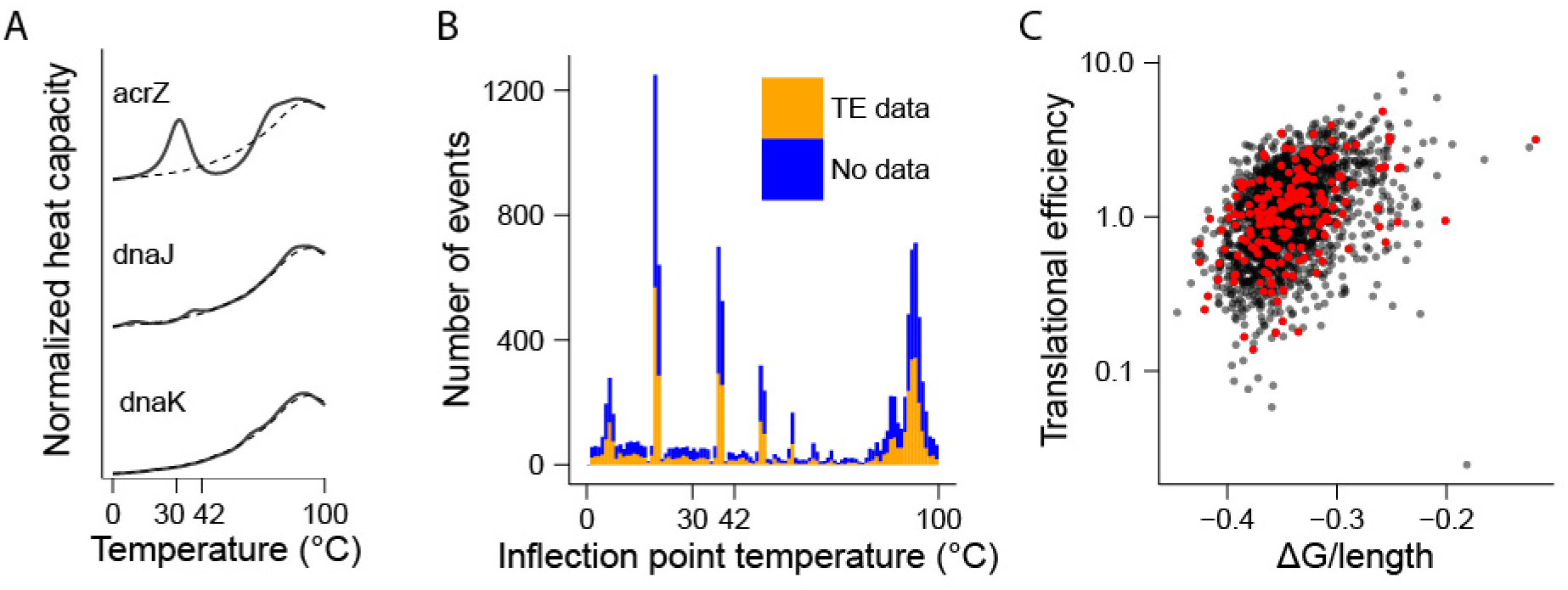
Few mRNAs have large predicted thermal melting transitions in the 30 °C to 42 °C range. (A) Example melting curves for genes with and without apparent transitions between 30 °C to 42 °C, calculated using the RNAheat program (solid lines). The average melting curve for all *E. coli* protein coding genes is shown for comparison (dashed lines). (B) Distribution of melting transition midpoint temperatures predicted by RNAheat. Transitions in genes for which TE data is available are shown in orange, others are shown in blue. (C) Correlation between TE and ORF-wide secondary structure (as in Fig 8B). Genes whose mRNAs have a large transition between 30 °C and 42 °C are highlighted in red.

*A minimal linear model can predict trends in translation efficiency*—Although no one factor can reliably predict translation efficiency, it is possible that similar combinations of sequence- or gene-specific factors control TE of subsets of genes (14). We used parameters derived from gene’s sequences and their consensus locations to fit the 30 °C log-transformed TE data to a linear model. The parameters used were the gene’s start codon; GC content; protein location; coding sequence length; genome position; predicted RBS binding strength; folding free energy of the 5’ UTR, 5’ end of the ORF and the combination of those regions; folding free energy divided by length of the whole ORF; and the calculated CAI and tAI for the full protein and its N-terminal 40 residues. Parameters and errors for the model are in Table S2. The model’s predictions correlate with the input data with an adjusted R² of 0.38, much better than the best individual parameter, CAI. However, relatively few parameters have a significant impact on the performance of the model, and a simpler model that includes only GC content, tAI, the predicted mRNA folding free energy per nucleotide, the presence of an ATG start codon and whether a protein is predicted to localize to the inner membrane, correlated with its input data with an adjusted R² of 0.33. FP levels predicted by the model from the measured mRNA levels correlated with measured FP levels with an R^2^ of 0.88, an improvement on the correlation between measured mRNA and FP levels (R^2^ = 0.81). Parameters for the model are in Table 2. Fig. 10 shows predicted versus measured TE values and the distributions of residuals of a prediction of footprint levels from the model, compared to that from the mRNA correlation. These parameters can all be calculated or predicted from an organism’s genome sequence. Therefore, this model may have some utility in predicting TE — and therefore predicting protein levels from transcriptome data — in other organisms with similar transcription machinery to *E. coli*.

**Table 2:** Parameters for reduced linear model. GC, G/C content (%); tAI, tRNA adaptation index; locIM, inner membrane localization (logical); startATG, presence of an ATG start codon (logical); mfePerNuc, RNA stability per nucleotide for entire ORF (kcal/mol/nuc).

| Parameter | Estimate | Standard error | Mean square error | F-statistic | P-value | log<sub>10</sub> P-value |
| --- | --- | --- | --- | --- | --- | --- |
| Offset | -0.852 | 0.162 |  |  |  |  |
| GC | 5.49 | 0.505 | 63.0 | 213 | $5.44 \times 10^{-47}$ | -46.3 |
| tAI | 7.21 | 0.402 | 249 | 840 | $5.97 \times 10^{-168}$ | -167.2 |
| locIM | -0.307 | 0.0243 | 89.0 | 301 | $5.17 \times 10^{-65}$ | -64.3 |
| startATG | 0.192 | 0.0283 | 19.1 | 64.6 | $1.18 \times 10^{-15}$ | -14.9 |
| mfePerNuc | 10.9 | 0.461 | 167 | 563 | $1.15 \times 10^{-116}$ | -115.9 |

**Fig. 10:**
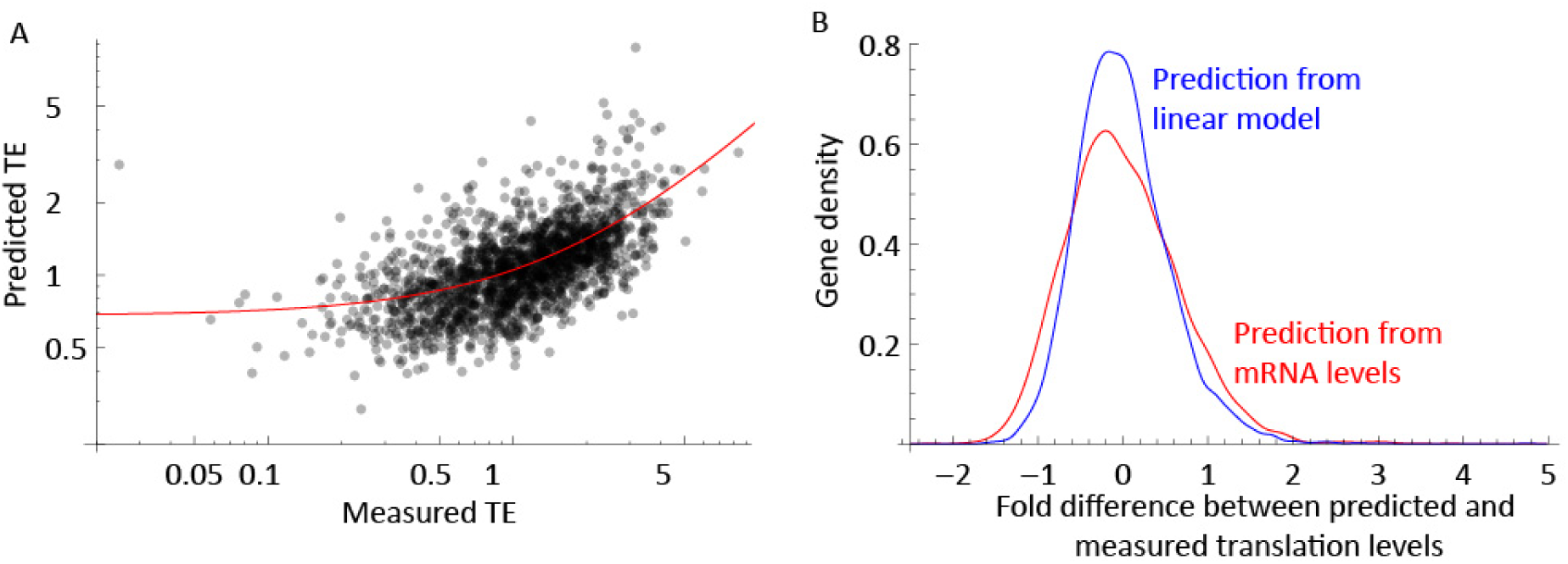
A linear model with five gene-specific parameters can explain 33% of the variation in translation efficiency. Translation data from three replicates at 30 °C was used to create a linear model to predict translational efficiency from gene sequence-specific data. (A) Predicted versus measured TE values. Each point represents a single measurement for a gene. The red line shows a linear regression fit of the data. (B) The fold-difference between the predicted and measured footprint CPKM values are plotted as smoothed density histograms. For comparison, the residuals for predicted footprint levels based on a linear fit to the mRNA levels are shown in red. Curves are scaled so that their integrals are equal. The narrower distribution of the linear model curve represents a better fit to the measured data.

## DISCUSSION

We have used ribosome profiling to measure transcription and translation in *E. coli* at 30 °C and after 10 and 20 minutes of heat shock at 42 °C. Translation rates are strongly dependent on mRNA levels, but the ratio of footprint to mRNA reads, TE, varies by over 100-fold across the genome (Fig. 1). Since translation initiation is thought to be governed by RNA hybridization, we expected to observe widespread changes in TE at different temperatures. However, both the overall TE (Fig. 2) and the patterns of ribosomal footprints (Fig. 5) were correlated at 30 °C and 42 °C, despite large changes in mRNA levels between conditions (Fig. 3 and 4). A gene’s TE is correlated with the overall structure of its mRNA (Fig. 8), which for most genes is predicted to be maintained over the temperature range investigated here (Fig. 9). Inner membrane proteins tend to be translated at a lower rate than other proteins (Fig. 8). A simple linear model can predict a third of the variation between genes’ translation (Fig. 10). This information is needed to incorporate heat shock into computational models of protein homeostasis such as FoldEco (3), and may be of use in predicting protein levels in other organisms.

This work uses a mild heat shock protocol, which is insufficient to activate *E. coli*’s known RNA thermometers, although it does result in increased transcription of σ^H^ regulon genes (Figs. 3 and 6) and increased translation of the *ssrA* ORF (Fig. 7). It is possible that transient changes in translation occur before the 10 minute time point measured here, since the transcriptional effects of the heat shock response peak around 5 minutes after a temperature shift (9). It is likely that higher temperatures cause greater changes in translation. However, translation efficiency appears to be very robust to the temperature changes used here. Recent observations by Bartholomäus and coworkers shows that *E. coli* subjected to a temperature jump from 37 °C to 47 °C do show translational changes in 129 genes, including members of the σ^H^ regulon (40). Genes whose translation increased upon heat shock had weak mRNA secondary structure content in their 5’ regions, suggesting that melting of these structures at high temperature may contribute to increased translation. TE of *ibpA* increased at 47 °C but TE of *rpoH* did not (40), supporting our observation that activation of σ^H^ is driven by transcriptional and post-translational mechanisms. Our RNA melting calculations (Fig. 9B) do not show widespread changes in structure in this temperature range. Translational changes are observed in heat-stressed mammalian cells over long time periods (41, 42), and several stress responses lead to translational attenuation mediated by eIF-2α phosphorylation (43). However, the acute effects of heat shock, mediated by RNA thermometers (44), are apparently not widespread in *E. coli*.

*Translation of inner membrane proteins*—The observation that inner membrane proteins and SRP client proteins (Fig. 8) are translated less efficiently than other proteins may be due to the effects of cotranslational translocation into the inner membrane via the SRP pathway. Translocation is faster than translation, and therefore unlikely to be rate-limiting (45), although footprint counts show that at least 10 times more ribosomes than SecYEG translocons are synthesized under these conditions. The geometry and steric constraints of the inner membrane may place a limit on the number of ribosomes that can simultaneously translate a particular gene. Polysomes in solution adopt a helical conformation that minimizes the space between ribosomes while maximizing the separation between emerging nascent chains (46). On a planar membrane surface, however, the density of ribosomes on an mRNA molecule may be limited by the packing of translocons and the requirement that the nascent chains are oriented in the same direction. Whatever the mechanism, it makes intuitive sense that nascent membrane proteins are kept away from each other to avoid their aggregation. *E. coli* mRNA molecules which encode membrane proteins have been shown to segregate to the membrane independently of translation (47), and the (as yet undetermined) factors responsible for this could also be involved in reducing ribosome density at the membrane.

*Predicting translation from sequence*—The coding and non-coding regions of mRNA can affect translation by several known mechanisms. mRNA structure and complementarity to the ribosome anti-Shine-Dalgarno sequence can affect both translation initiation and elongation (27, 48). The genetic code is degenerate — most amino acids are encoded by several codons, but organisms use them selectively (34, 49). Codon use can alter the translational elongation rate as ribosomes wait for more or less abundant aminoacyl tRNAs (34, 49–51), but different mRNA sequences also have different structural propensities which influence TE (50, 52–54). The 5’ end of open reading frames seems to be particularly important in determining how a gene is translated (50, 55, 56). The relative contribution of these effects is the subject of much research and debate (16, 57). Natural genomes have evolved under constraints that are not fully understood, and there may be as-yet unidentified mechanisms that control TE. Perhaps the roles of RNA binding proteins and chaperones, or ribosomal protein S1 (58) in *E. coli* are more significant than is currently appreciated. From the point of view of synthetic biology, technology may have overtaken theory: Kosuri et al. suggest that screening large libraries of synthetic constructs may be a more effective method than rational design for finding genes with the desired expression properties (59). However, optimization of RBS sequences is an effective way to modulate the expression of a particular gene (60).

Ribosome profiling approaches have shown a clear correlation between ORF-wide mRNA structure and TE across the *E. coli* genome (20), but not with other sequence-dependent parameters (12, 14). Our analysis supports this, suggesting that factors affecting elongation rather than initiation — ORF-wide structure (Fig. 8A), tAI (Fig. 8B) and cotranslational translocation (Fig. 8E) — determine TE. This is despite the “snapshot” nature of a ribosome profiling measurement: an increased elongation rate would reduce the time a ribosome spends on an mRNA molecule, thereby reducing apparent ribosome density. The work presented here shows that control of TE is robustly maintained against temperature changes that might be expected to differentially alter RNA:RNA interactions. Prediction of the temperature dependence of mRNA structure indicates that there are relatively few structural transitions in the 30 °C to 42 °C range (Fig 9), which supports the hypothesis that *E. coli* has evolved to minimize temperature-dependent translational changes. Prediction of long RNA structures is challenging. The simple approach taken here has less predictive ability than the measurement of RNA structures *in vivo*. Recent technical advances in such measurements (20, 61) should lead to better structure prediction algorithms. However, the ability of the model presented here to predict TE from sequence-dependent parameters could be useful for interpreting existing or new transcriptome data, whether from *E. coli*, other bacteria or metagenomics studies. Because the necessary parameters are easily calculated or predicted from sequences, this approach suggests a way to refine estimates of protein levels from only a genome sequence. It remains to be seen how widespread these patterns of translational control are, both within *E. coli* strains and in other organisms.

## EXPERIMENTAL PROCEDURES

The procedure for ribosome footprinting and cDNA library preparation was modified slightly from that published in Oh et al. (5). A detailed protocol is given in (62).

*Growth conditions—E. coli* K12 MG1655 cells were grown in 200 ml cultures of EZ MOPS defined rich media (Teknova) at 30 °C, 200 rpm. A total of 18 cultures were grown in the batches shown in Table 1, starting from fresh overnight cultures of the same glycerol stocks of bacteria. For the temperature shift experiments, cultures of untransformed cells were grown to OD_600_ of 1 in media without antibiotic at 30 °C, then diluted 3-fold into fresh media at either 30 °C or 42 °C, grown for 10 or 20 minutes then harvested at an OD_600_ of between 0.4 and 0.5. We used shaking waterbaths to stabilize the media temperature. For σ^H^ expression, cultures were transformed with a pBAD vector containing either wild-type or I54N σ^H^, or with an empty pBAD vector. Cultures were grown in media containing 0.1 mg/ml ampicillin to an OD_600_ of 0.2, induced with 0.2% arabinose, grown for a further 20 minutes and then harvested at an OD_600_ of between 0.4 and 0.5.

*Harvesting and lysis*—Cells were harvested by rapid vacuum filtration through a 90 mm diameter 0.2 μm pore filter, scraped off the filter with a scoopula then immediately plunged into a 50 ml tube full of liquid nitrogen. Harvesting time from decanting to freezing was between 90 and 120 s. The frozen cells were scraped off the scoopula into the bottom of the tube. Nitrogen was allowed to evaporate at -80 °C and cell pellets were stored frozen at -80 °C until lysis, typically overnight. Frozen cell pellets were lysed at liquid nitrogen temperatures using a bead beater with steel tubes, silicone caps and a single 5 mm steel ball (BioSpec). Frozen cell pellets were decanted into pre-cooled tubes containing 600 μl of frozen lysis buffer (100 mM NH_4_Cl, 10 mM MgCl_2_, 5 mM CaCl_2_, 20 mM Tris-Cl pH 8, 0.1% NP-40, 0.4% Triton X-100, 50 μg/ml chloramphenicol, 100U/ml DNase I). All components were RNase-free. Chloramphenicol in the lysis buffer stalled the ribosomes on mRNA to allow for analysis of ribosomal pausing patterns. A pre-cooled ball was added, the tubes were capped, and shaken at full speed on a Mini-Beadbeater-1 for 6 cycles of 10 seconds, and were re-cooled in liquid nitrogen for 45s – 1 min between cycles. This treatment gave satisfactory lysis of the cells without apparent thawing; increasing the number of cycles ran the risk of splitting the silicone caps.

*Ribosome footprinting and polysome profiling—*Cell lysate was thawed in the steel tubes at 30 °C, incubated on ice for 10 minutes, then transferred into 1.7 ml Eppendorf LoBind DNA tubes and centrifuged at 14,000 rpm in a microfuge for 10 minutes at 4 °C. The supernatant was removed and a 30 μl aliquot was snap frozen and stored at -80 °C; this sample was used for total cellular RNA. Two 180 μl aliquots of the remaining lysate were used for polysome profiling and ribosome footprinting. One aliquot was incubated with streptococcal nuclease S7 (Roche) for one hour at 25 °C to produce ribosome-protected footprints; the other was incubated without the nuclease as a control to confirm that cells were translating and that the nuclease treatment had digested the mRNA to leave monosomes. The nuclease digestion was quenched with EDTA and the lysates cooled on ice. The lysates were immediately loaded onto a 10-40% sucrose gradient (in buffer containing 100 mM NH_4_Cl, 10 mM MgCl_2_, 5 mM CaCl_2_, 20 mM Tris-Cl pH 8, 100 mM chloramphenicol and 2 mM dithiothreitol) in Beckman centrifuge tubes and centrifuged at 35,000 rpm for 2.5 hours in a Beckman SW-41 rotor. Polysome profiles were measured by pushing the gradients out of the tube with 60% sucrose solution and monitoring RNA absorbance at 260 nm. The fractions corresponding to the center of the monosome peak were collected for the nuclease-digested samples, pooled and frozen at -80 °C.

*RNA extraction*—RNA was extracted from the total RNA and monosome fractions by the hot acid phenol chloroform method. RNA was precipitated with isopropanol after adding sodium acetate and GlycoBlue (Thermo Fisher) as a coprecipitant. Total RNA was enriched for mRNA by purification with MegaClear and RiboMinus kits to remove small RNAs and rRNA respectively, following the kit manufacturer’s instructions. The remaining RNA was fragmented by incubation at 95 °C, pH 9.3 in sodium carbonate buffer for 40 minutes. Monosome fractions and fragmented mRNA were loaded onto a 15% polyacrylamide TBE-urea gel (Thermo Fisher) and run at 200V for 1 hour. Gels were stained with SYBR-Gold (Thermo Fisher) and the band corresponding to footprint-sized oligonucleotides was excised. RNA was extracted from the gel by crushing the gel slice and shaking at 70 °C for 10 minutes. Gel fragments were removed by filtration and the RNA was and precipitated as above.

*cDNA sequencing library preparation—*The extracted RNA footprints and fragments were dephosphorylated by incubating with T4 polynucleotide kinase (NEB) without ATP. RNA was ligated to a 13-nucleotide adaptor (CTGTAGGCACCATCAAT) (IDT) using truncated T4 RNA ligase. The ligated products were purified using a Zymo RNA cleanup column which removed unligated adaptor and concentrated the RNA. Ligated RNA was reverse transcribed by Superscript III (Thermo Fisher) using a primer complementary to the adaptor sequence, which also contained the sequences necessary for PCR amplification separated by a peptide spacer (5Phos/GATCGTCGGACTGTAGAACTCTGAACCTGTCGGTGGTCGCCGTATCATT/iSp18/CA CTCA/iSp18/CAAGCAGAAGACGGCATACGA ATTGATGGTGCCTACAG where iSp18 is the peptide spacer). Full-length cDNA was gel-purified as before from a 10% polyacrylamide TBE-urea gel. RNA was hydrolysed at pH 14, 95 °C for 40 minutes, leaving single-stranded cDNA. cDNA was circularised using CircLigase (Epicentre) to produce single-stranded, circular DNA molecules which included the two complementary sequences for PCR amplification needed to make Illumina sequencing libraries. Footprint libraries tend to be contaminated with specific rRNA sequences, which were removed by hybridization with specific biotinylated oligonucleotides (IDT) followed by capture on streptavidin magnetic beads (Thermo Fisher). Circular cDNA libraries were amplified by 8-12 cycles of PCR with Phusion polymerase (NEB). Primers encoded an Illumina TruSeq barcode sequence at the 5’ end of the insert to allow for multiplexing. Amplified libraries were gel-purified on 8% polyacrylamide TBE gels; the sample that had experienced the largest number of cycles without showing large overamplification products was excised from the gel and extracted overnight at 20 °C.

*Sequencing and alignment*—Libraries were quantified by Agilent BioAnalyzer, and pooled to give a final sequencing library containing 12 barcoded samples. Libraries were sequenced on an Illumina HiSeq 2500 at the TSRI sequencing core facility. Each barcoded sample typically gave 10-20 million single-end, 100bp reads. Base calling and demultiplexing were done with Illumina’s Casava software. Adaptor sequences were removed from reads using the fastx_clipper application http://cancan.cshl.edu/labmembers/gordon/fastx_toolkit to leave footprint or fragment sequences. These were aligned to the *E. coli* K12 MG1655 genome (NC_000913) using Bowtie (63). Most samples resulted in between one and three million mapped reads; the unmapped reads were mostly due to contaminating rRNA, which was not removed from our libraries as successfully as was reported previously. The number of reads mapped to a particular nucleotide was counted using an in-house Python script (modified from that used in Oh et al. (5)) that averaged each read over the central nucleotides in the sequence. Since footprint read lengths are non-uniform in bacteria, the exact position of the ribosome peptidyl transferase site on each read cannot be precisely determined. The resulting .wig files were processed with in-house R scripts using a variety of analysis packages as described below.

*Calculation of per-gene CPKM values and translational efficiencies*—Reads were mapped to a list of protein-coding gene positions taken from EcoCyc (64). RNA genes, pseudogenes and phage Ins elements were excluded from the list. Reads mapping to pairs of genes with similar sequences (*tufA* and *tufB*; *gadA* and *gadB*; *ynaE* and *ydfK*; *ldrA* and *ldrC*; *ybfD* and *yhhI*; *tfaR* and *tfaQ*; *rzoD* and *rzoR*; and *pinR* and *pinQ*) were aligned randomly to one homolog and the total counts used for determining mRNA and footprint levels, but were excluded from further analysis. A meta-analysis of reads mapped to well-translated genes (those with at least 128 footprint counts) showed a similar enrichment of ribosome density at the 5’ end of genes. For each footprint dataset, we corrected for this bias by dividing the read count at each codon within a gene by the normalized average ribosome density at that position from well-expressed genes (12). Read counts per gene were calculated from either the raw reads (for mRNA counts) or normalized reads (for footprints). We calculated counts per kilobase million (CPKM) for each gene in each experiment using the EdgeR package (23), which normalizes the counts per gene by the total number of aligned reads, then corrected for gene length. These CPKM values are the basic measurement of a gene’s mRNA level and ribosome density, and can be directly compared between datasets. Translation efficiency (footprint CPKM to mRNA CPKM) ratios were calculated pairwise for each gene in each experiment where that gene had at least 64 raw (unnormalized) counts for both mRNA and footprints. The mean TE for each condition was used to assess the influence of gene-specific parameters on TE.

*Gene-specific parameters*—Sequences and locations were taken from EcoCyc (64). We consider only protein-coding genes in this analysis, excluding genes with close homologs as above, selenoproteins, and proteins with frameshifts or stalling sites (*fdhF*, *fdoG*, *fdnG*, *prfB*, *dnaX*, *secM*, and *tnaC*). RNA secondary structures were calculated with Vienna RNA (39). We used the RNAfold program to determine minimal free energy values for every protein coding ORF, and the RNAheat program to calculate melting curves. GC content, codon adaptation index and tRNA adaptation index scores for each gene were calculated with the cai function from the seqinr R package (65) using data from (49) and (34). Protein locations were taken from the consensus data in Diaz-Mejia et al (36). Ribosome binding site calculations at different temperatures were calculated using the RBS Calculator software (66).

*Linear modeling*—Linear models for translation efficiency were calculated using Wolfram Mathematica. Mean translation efficiency was fitted to a model incorporating all of the parameters calculated for each gene (Table S2). A second model was calculated using only five parameters which contributed most to the fit of the first model (Table 2). We used the model to predict the ribosome footprint CPKMs for each gene given that gene’s mRNA CPKM. The correlation between the predicted and measured footprint CPKMs is reported. To visualize the increased information content of the model, we compared the distribution of residuals from the fit of the model to the footprint counts with the residuals calculated from a simple linear regression of the mRNA and footprint CPKMs.

*Caulobacter crescentus data*—Data for *C. crescentus* in rich (PYE) media was taken from Schrader et al. (38). Homologous genes were identified using the EcoCyc database. Protein localizations were from annotations in the BioCyc *C. crescentus* NA1000 database. The inner membrane class in Fig. 8F is a combination of the single- and multi-pass membrane proteins defined by BioCyc; peripheral membrane proteins were assigned to the cytosol

## ACKNOWLEDGEMENTS

This work was supported by NIGMS grant GM101644. Sequencing data are deposited in GEO under accession GSE90056. We thank Dr. Jonathan Weismann, Dr. Carol Gross, Dr. Gene-Wei Li and members of their laboratories for assistance with ribosome profiling experiments; Dr. Jamie Williamson for assistance with sucrose gradients; Dr. Steven Head and the TSRI sequencing core for assistance with sequencing; Ryan Thomson for assistance with data analysis; and Dr. Howard Salis for the ribosome binding site calculations. We thank all of these individuals and members of the Kelly laboratory for helpful discussions and critical reading of the manuscript.

### CONFLICT OF INTEREST

The authors declare that they have no conflicts of interest with the contents of this article.

### AUTHOR CONTRIBUTIONS

GJM, JWK and ETP conceived and designed the study. GJM acquired and analyzed the data with assistance from DHB for the ribosome profiling experiments. GJM wrote the paper with input from all authors. DHB shared unpublished data relating to the control of translation that was critical to the interpretation of the results. All authors reviewed the results and approved the final version of the manuscript.

## SUPPLEMENTAL TABLE LEGENDS

**Table S1:** Excel spreadsheet of raw reads and EdgeR-calculated CPKM values for all genes under all conditions. Gene data is taken from the EcoCy database (64). Only protein coding genes are included in the CPKM analysis. Data for homologous genes was summed and is labeled as “geneA.geneB” in the CPKM data. One replicate of the 42 °C, 20 minute time point had anomalous data and was excluded from further analysis. A plot of mRNA CPKM vs footprint CPKM for the three replicates of this condition shows the similarity of two (red and blue) datasets compared to the other (green).

**Table S2:** A linear model using 25 gene-specific parameters was used to predict TE values for genes. Parameter estimates and goodness-of-fit information is shown for comparison to the minimal linear model described in Table 2.

